# Covert attention for uncertainty reduction during sequential inference

**DOI:** 10.1101/2023.11.28.569036

**Authors:** F. Javier Domínguez-Zamora, Guillermo Horga, Jacqueline P. Gottlieb

## Abstract

A major question in attention research is how the brain identifies task-relevant stimuli in the absence of exogenous instructions or cues. Recent studies propose that endogenous attention control expected information gains (EIG) or, equivalently, minimizes decision uncertainty, but the mechanisms of this process are not understood. We show that, in a task in which participants covertly attended to decision-relevant stimuli, their perceptual sensitivity (d’) for discriminating the stimuli depended on the diagnosticity of the stimuli and the participants’ prior decision uncertainty, consistent with Bayesian EIG. The fronto-parietal network, in particular left areas V3A/B and IPS1/2, integrated uncertainty with diagnosticity in a manner correlating with behavioral effects on d’, and uncertainty signals relied on interactions between this network and the medial prefrontal cortex (mPFC). The findings show that covert attention can be deployed based on EIG and reveal the neural mechanisms of this process.

## INTRODUCTION

Natural environments are replete with sensory information, but only a small subset of that information is relevant for learning or actions. Thus, making adaptive decisions requires individuals not only to choose appropriate actions but to appropriately orient their attention – i.e., identify and focus on task relevant stimuli while avoiding irrelevant and time-consuming distractors.

Converging evidence shows that, in humans and monkeys, the task-related control of attention depends on visual and fronto-parietal networks^1-3^. In monkeys, neurons in the frontal eye field and lateral intraparietal area selectively encode the locations of presented or anticipated task-relevant stimuli^2,3^. These neurons predict not only the goal of a saccadic eye movement (an “overt” orienting response) but also covert shifts of attention – i.e., the internal prioritization of a specific stimulus manifested as an improvement in sensory discriminability (d’)^2,3^. Consistent with these findings in monkeys, human areas V3A/B, the intraparietal sulcus (IPS) and the frontal eye field (FEF) encode an anticipatory attention orienting response – i.e., the selection of a location where a task-relevant stimulus is expected to be ^4-7^. Thus, a network comprised of visual and fronto-parietal areas seems to mediate the advance selection of information sources (or channels), to enhance the discriminability of specific stimuli that latter appear in those channels.

A longstanding open question, however, is how visual and fronto-parietal areas identify relevant stimuli. How does the brain “know” which information sources are relevant for a task^8,9^? Converging evidence suggests that this process involves not only the fronto-parietal network but also the executive network, in particular the medial prefrontal cortex (mPFC)^10-12^, but it is unclear whether or how the mPFC is related to perceptual discriminability.

The Bayesian framework of probabilistic reasoning postulates that, since the key role of attention is gathering information -- or, equivalently, reducing uncertainty – optimal attention control is based on expected information gains (EIG): the extent to which a stimulus, if discriminated in detail, will reduce uncertainty about future task-relevant actions or states. Moreover, Bayesian logic specifies that EIG requires the integration of diagnosticity and prior uncertainty. Diagnosticity measures evidence strength, and is the probability that a stimulus, if discriminated in detail, will correctly predict future actions or states. Prior uncertainty is the extent to which the decision maker is initially unsure about the correct choice and needs information from additional stimuli.

As a simple example, consider a driver who reaches an intersection and decides if to step on the brake or the gas. If the driver attends to the traffic light and receives an unambiguous signal (e.g., red) his decision uncertainty is quickly resolved, obviating the need to attend to additional stimuli. However, if the driver sees an ambiguous yellow, his uncertainty remains high, incentivizing him to attend to additional stimuli (e.g., the pedestrians or the speed of the traffic). Thus, as evidence accumulates preceding a choice, attention should be flexibly allocated in response to changes in decision uncertainty. However, it is unknown if perceptual discriminability shows such uncertainty-based control or how this may be mediated by the executive and fronto-parietal networks.

To examine this question, we used fMRI imaging during a task in which participants covertly attended to briefly-flashed, near-threshold stimuli that were relevant to a binary choice. The stimuli conveyed probabilistic information that had high or low diagnosticity (hiD or loD) – i.e., predicted the correct choice with high or low accuracy – and could appear in conditions of high or low decision uncertainty. Moreover, the stimuli could flash in the right or left hemifield and, during a delay period preceding the flash, participants were informed about the locations in which hiD and loD stimuli were expected to flash.

We show that, consistent with Bayesian EIG, participants had higher perceptual sensitivity (d’) for discriminating the stimulus at a hiD relative to loD location and, crucially, this spatial bias was stronger under high versus low uncertainty. Multivariate decoding analyses showed that the anticipated location of the hiD stimulus (right vs left hemifield) was decoded with above chance accuracy from the bilateral fronto-parietal network and, crucially, this decoding was enhanced by uncertainty. The effect of uncertainty was particularly prominent in left areas V3A/B and IPS1/2 and, in the latter, correlated with the effect of uncertainty in individual participants’ d’. Finally, psychophysiological interaction analyses (PPI) showed that uncertainty enhanced the functional connectivity between the mPFC and left V3A/B, suggesting that uncertainty information may reach the fronto-parietal network through interactions with the mPFC. The findings reveal interactions between the executive and fronto-parietal networks that can mediate EIG-based adjustments in perceptual sensitivity.

## RESULTS

### Task

Participants (n=30) performed a probabilistic inference task in which they discriminated near-threshold visual information relevant to a choice. Participants were told that each trial had a hidden jar that contained a mixture of upright and inverted letter “Ts”, such that the majority of the letters could be upright or inverted (majU or majI; Fig. 1A) and, at the end of the trial, they should report the letter majority orientation. To make this decision, participants maintained central fixation and viewed two letters that were randomly drawn from the jar. The 1^st^ letter was shown clearly at the center of gaze (Fig. 1B, “1^st^ letter”). After an anticipatory delay (Fig. 1B, “Anticipation”), the 2^nd^ letter flashed briefly at an eccentric location, requiring attention to discriminate its orientation (Fig. 1B, “2^nd^ letter”). Participants reported the 2^nd^ letter orientation immediately after the flash (Fig. 1B, “Orientation?”), after which they reported the majority orientation and received feedback regarding this choice (Fig. 1B, “Decision” and “Decision Feedback”). Fixation was enforced by eye tracking and, if participants broke fixation before the 2^nd^ letter flash, the trial was immediately terminated and repeated later on.

**Figure 1:**
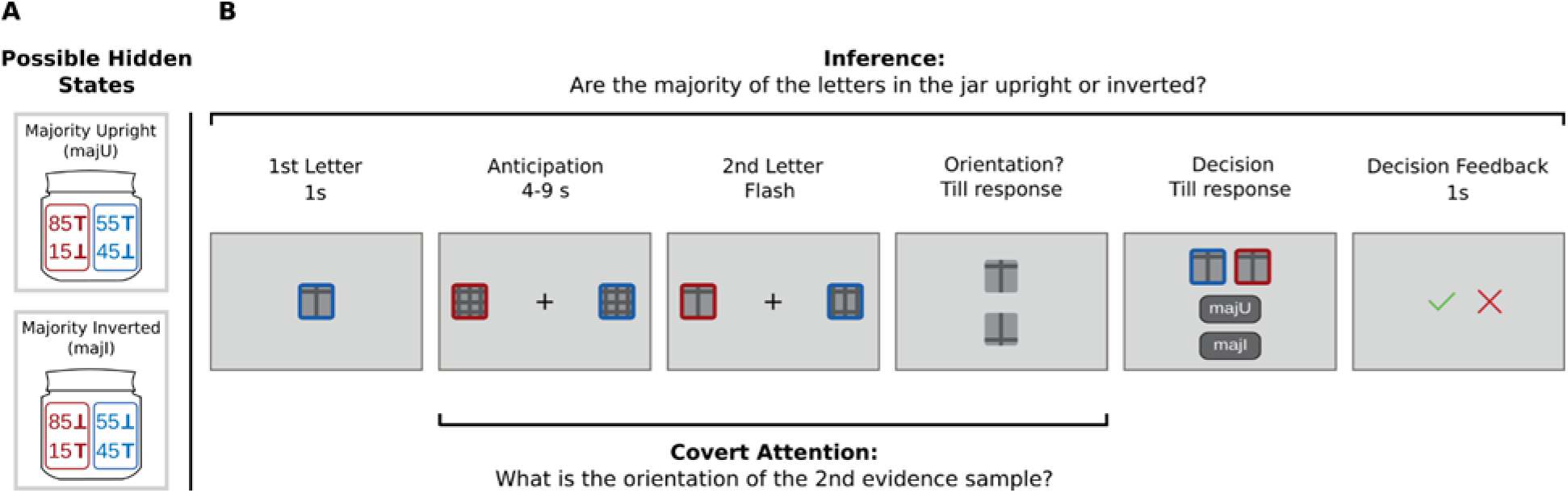
Task. **A. The hidden states.** Each trial had one of two possible hidden states, consisting of a jar with 200 hundred letter “Ts”, of which the majority could be upright (top, majU) or inverted (bottom, majI). Within each jar, the letters were divided into two boxes that had the same majority orientation but had, respectively, 55% or 85% of letters matching the majority orientation (respectively, loD and hiD). The boxes were identified by color (red vs blue, counterbalanced across participants); throughout the manuscript, we use red to indicate hiD for simplicity. **B. Trial events.** Participants viewed two samples of evidence – a 1^st^ letter followed by a 2^nd^ letter after a variable delay (“Anticipation”). The 1^st^ letter was presented at the center of gaze, and the 2^nd^ was flashed briefly at a peripheral location. Participants reported the orientation of the 2^nd^ letter immediately after the flash (“Orientation?”) and completed the trial by reporting the majority orientation of the jar (“Decision”) and receiving feedback about this report. While viewing the letters, participants were required to maintain central fixation, enforced by eye tracking. After the 1^st^ letter presentation, peripheral-colored boxes appeared at 8 degrees in the right and left visual field, marking the locations where the 2^nd^ letter would flash if it had, respectively, hiD or loD. Before making their final decision about the jar type, participants were reminded of the evidence they received, including the letters’ diagnosticities (colored frames), the true orientation of the 1^st^ letter and the orientation they reported for the 2^nd^ letter whether it was correct or erroneous.

Our focus was on how participants oriented attention during the anticipatory delay – i.e., when they prepared to discriminate the 2^nd^ letter orientation after viewing the 1^st^. To ensure that participants could estimate EIG in this epoch, we signaled if each letter had high or low diagnosticity (hiD or loD) – i.e., predicted the majority orientation with high or low accuracy – using a colored frame surrounding the letter. Specifically, participants were told that each letter could come from a “red” or “blue” box in which, respectively, 85% or 55% of the letters matched the majority orientation (Fig. 1A). Thus, if a 1^st^ letter came from the hiD box, it signaled the majority orientation with 85% accuracy whereas, if it came from the loD box (Fig. 1B) it had only 55% accuracy. (For simplicity, we refer to red and blue as indicating, respectively, hiD and loD but, during data collection, the color-diagnosticity mappings were counterbalanced across participants.)

The diagnosticities of the 1^st^ and 2^nd^ letters were independently randomized, with the 4 possible combinations of hiD/loD letters interleaved in a session. Because the letters were presented sequentially, the diagnosticity of the 1^st^ letter determined the decision uncertainty that participants had before viewing the 2^nd^. Thus, if the 1^st^ letter had hiD, participants had low uncertainty about the jar type (loU) while expecting the 2^nd^ letter flash; conversely, if the 1^st^ letter had loD, they had high uncertainty (hiU). In addition, two peripheral colored frames remained visible during the anticipatory delay indicating the location where the 2^nd^ letter would flash (in the right or left hemifields) if it had, respectively, hiD or loD (these locations were kept fixed across short interleaved trial blocks; see Methods and Fig. 1B, “Anticipation”). Thus, during the anticipatory delay, participants could combine spatial information about diagnosticity with their level of decision uncertainty to allocate attention to the possible flash locations based on their EIG.

Importantly, while participants were instructed and incentivized to maximize their decision accuracy, they received no instructions for a specific attention strategy. Individual trials ended with feedback on the accuracy of the final decision (Fig. 1B, “Decision Feedback”) and, at the end of the session, participants received a monetary bonus based on the accuracy of this choice (announced in advance; Methods). However, no rewards or feedback were given for the discrimination report, allowing participants to endogenously select the attentional policy they considered beneficial for the task.

### Behavior

Bayesian models prescribe that the EIG of the competing 2^nd^ letter locations will with both diagnosticity and prior uncertainty. To illustrate this prediction, we simulated a Bayesian agent who makes optimal inferences based on the information it has, but discriminates the 2^nd^ letter orientation with variable accuracy (see Methods). The results are shown in detail in the insets of Fig. 2A, which plot the agent’s decision accuracy as a function of its discrimination accuracy for the 4 letter combinations. If the 2^nd^ letter has loD, the traces are shallow: accurately discriminating the orientation of a loD 2^nd^ letter has little benefit for the final decision accuracy (Fig. 2A inset, left panel). In contrast, if the 2^nd^ letter has hiD, the slopes are much higher particularly under high uncertainty (Fig. 2A, inset, right, solid vs dashed traces). The slopes of these traces indicate EIG – i.e., the extent to which more accurate sensory discrimination enhances decision accuracy (reduces decision uncertainty). The main panel in Fig. 2A plots the 4 slopes and clearly depicts the characteristic uncertainty x diagnosticity interaction: EIG is higher for hiD versus loD 2^nd^ letters, and this difference increases with uncertainty.

**Figure 2:**
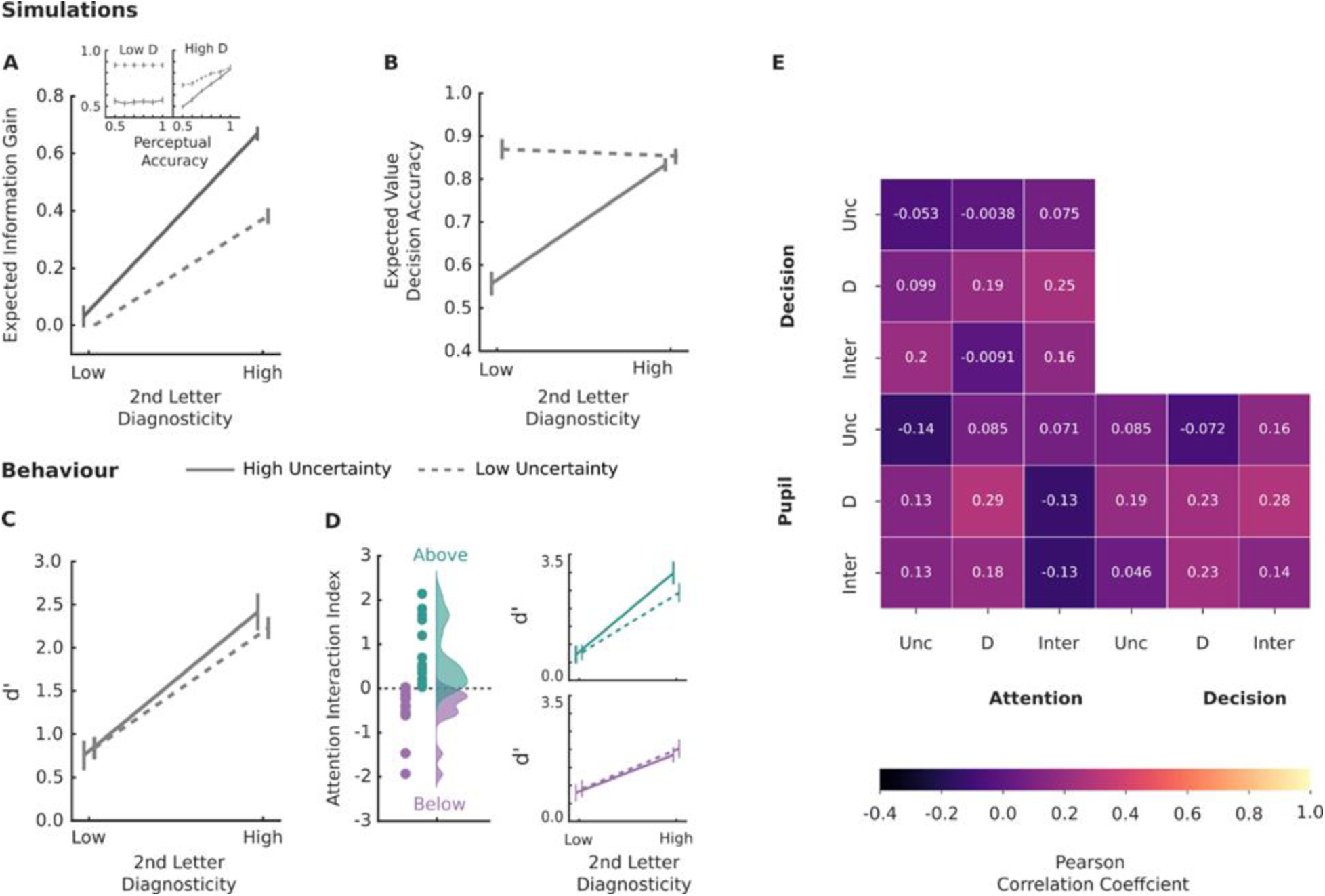
Integration of uncertainty and diagnosticity for attention control. **A. Simulated EIG** Inset: The decision accuracy (y-axis) as a function of perceptual accuracy (x-axis) for a simulated Bayesian agent. Left panel: loD 2^nd^ letter. Right panel: hiD 2^nd^ letter. Solid: high uncertainy trials; Dashed: low uncertainty trials. **Main panel:** EIG calculated as the slope at each uncertainty-diagnosticity combination in A. Each point shows mean and SEM over 30 model simulations. **B. Simulated expected value** for an agent with perfect perceptual accuracy for each trial type (mean and SEM over 30 simulations). **C. Perceptual discriminability** -- d’ -- in the same format as **A**. Each point shows the mean and SEM, N = 30 participants. **D**. **Inter-individual variability of attentional strategies.** The left panel shows indices indicating uncertainty-diagnosticity interactions in d’ for individual participants (circles) and their distributions (density histograms). The colors show the results of a median split, distinguishing between participants with above-median (green) and below median (purple) interactions. The right panel shows the d’ pattern for each group in the same format as **C**. **D. Independence of effects on d’, arousal and decisions** Correlation matrix between indices capturing the effects of diagnosticity (D), uncertainty (Unc) and interactions (Inter) measured for d’, pupil and decision weights (see Table 1 and **Supplementary Note 1** for details). The Pearson coefficients are shown by the color scale and white numbers. There were no statistically significant correlations.

**Table 1:** Diagnosticity, Uncertainty and Interaction indices for different behaviors. All values show mean and SE across N = 30 participants. For all measures (rows), the diagnosticity index is the difference in d’ for hiD and loD 2^nd^ letter discrimination divided by the sum; the uncertainty index is the difference in d’ between high and low uncertainty trials divided by the sum; and the interaction index is the difference between the diagnosticity indices on hiU vs loU trials. P values are from two-tailed Wilcoxon tests against 0.

|  | Diagnosticity |  |  | Uncertainty |  |  | Interaction |  |  |
| --- | --- | --- | --- | --- | --- | --- | --- | --- | --- |
|  | Mean | SE | p | Mean | SE | p | Mean | SE | p |
| **d'** | **1.218** | 0.227 | **$1.360 \times 10^{-5}$** | 0.054 | 0.084 | 0.734 | 0.160 | 0.161 | 0.465 |
| **Pupil size** | $1.109 \times 10^{-4}$ | 0.002 | 0.959 | **0.009** | 0.004 | 0.037 | 0.507 | 0.465 | 0.349 |
| **Decision Weights** | **1.428** | 0.130 | **$2.353 \times 10^{-6}$** | **1.276** | 0.149 | **$5.383 \times 10^{-6}$** | **1.719** | 0.326 | 0.000 |

Importantly, the EIG pattern is distinct from a trial’s expected value (EV; Fig. 2B). EV depends on the maximal possible decision accuracy (i.e., the total information available) and is high if at least one letter has hiD. In contrast, EIG indexes the marginal accuracy benefit due specifically to the 2^nd^ letter discrimination, over and above the information conveyed by the 1^st^ (Fig. 2A).

To test if participants oriented attention based on EIG, we focused on the discrimination report and calculated d’ -- a criterion-free measure of perceptual sensitivity that is the gold-standard for detecting covert attentional modulations. Consistent with EIG, the average d’ was higher for hiD vs loD 2^nd^ letters particularly under higher uncertainty (Fig. 2C). However, while the effect of diagnosticity was robust, the effects of uncertainty were not significant in the group-level analysis (2-way ANOVA, diagnosticity main effect, p < 10-4; uncertainty main effect: p = 0.78; interaction: p = 0.463, N = 30 participants).

We hypothesized that, because participants were not instructed or trained on how to attend, they may have pursued individually heterogeneous strategies. To examine this hypothesis, we measured the effects of diagnosticity and uncertainty on each participant’s data and, crucially, measured the diagnosticity x uncertainty interaction as the difference between the effects of diagnosticity on hiU vs loU trials (see Table 1 for a full explanation and descriptive statistics). As shown in Fig. 2D, 17/30 participants showed positive interactions and only 2/30 showed strongly negative interactions. Thus, many participants tended to enhance the spatial bias based on uncertainty, consistent with the relative EIG of the competing locations. Other participants, in contrast, pursued an uncertainty-independent strategy of consistently attending to the hiD location regardless of uncertainty (no interactions; see Discussion for possible explanations of these strategies). To capture this variability, we divided participants into two equal sized groups with interaction indices above- and below- the median value (Fig. 2D, green vs purple), whose interaction indices differed significantly (mean (SEM) index of, respectively, 0.77 (0.71) vs -0.45 (0.55); p < 10-3, t=4.6, N = 15 participants in each group).

### Left IPS integrates uncertainty with diagnosticity dependent spatial bias

To understand the neural mechanisms of EIG-based attentional orienting, we analyzed BOLD responses during the anticipatory delay (400-800 ms after 1^st^ letter onset). We used the probabilistic atlas of Wang et al.^13^ to independently identify 5 regions of interest (ROI) in the fronto-parietal network – namely, the frontal eye field (FEF) and areas V3A, V3B, IPS1 and IPS2 in both hemispheres (Fig. 3A). In a first step, we trained support vector machines (SVM) to decode the expected location of the hiD 2^nd^ letter (i.e., whether the hiD peripheral marker was in the right vs left hemifield) regardless of uncertainty (Methods, Multivariate Analysis). Consistent with their documented response to spatial anticipatory attention^4-7^ all the fronto-parietal ROIs showed above-chance decoding accuracy for the hiD 2^nd^ letter location: (mean (SEM): Left V3A=0.59 (0.003); Left V3B=0.6 (0.003); Left IPS1=0.56 (0.004); Left IPS2=0.55 (0.003); Left FEF=0.54 (0.003); Right V3A=0.6 (0.004); Right V3B=0.6 (0.004); Right IPS1=0.55 (0.003); Right IPS2=0.53 (0.003); Right FEF= 0.54 (0.003); all p’s < 0.05, permutation test). Thus, complementing the univariate analyses in previous studies^4-7^, advance spatial orienting to the hiD location was reliably decoded from the multivoxel response pattern in attentional ROIs.

**Figure 3:**
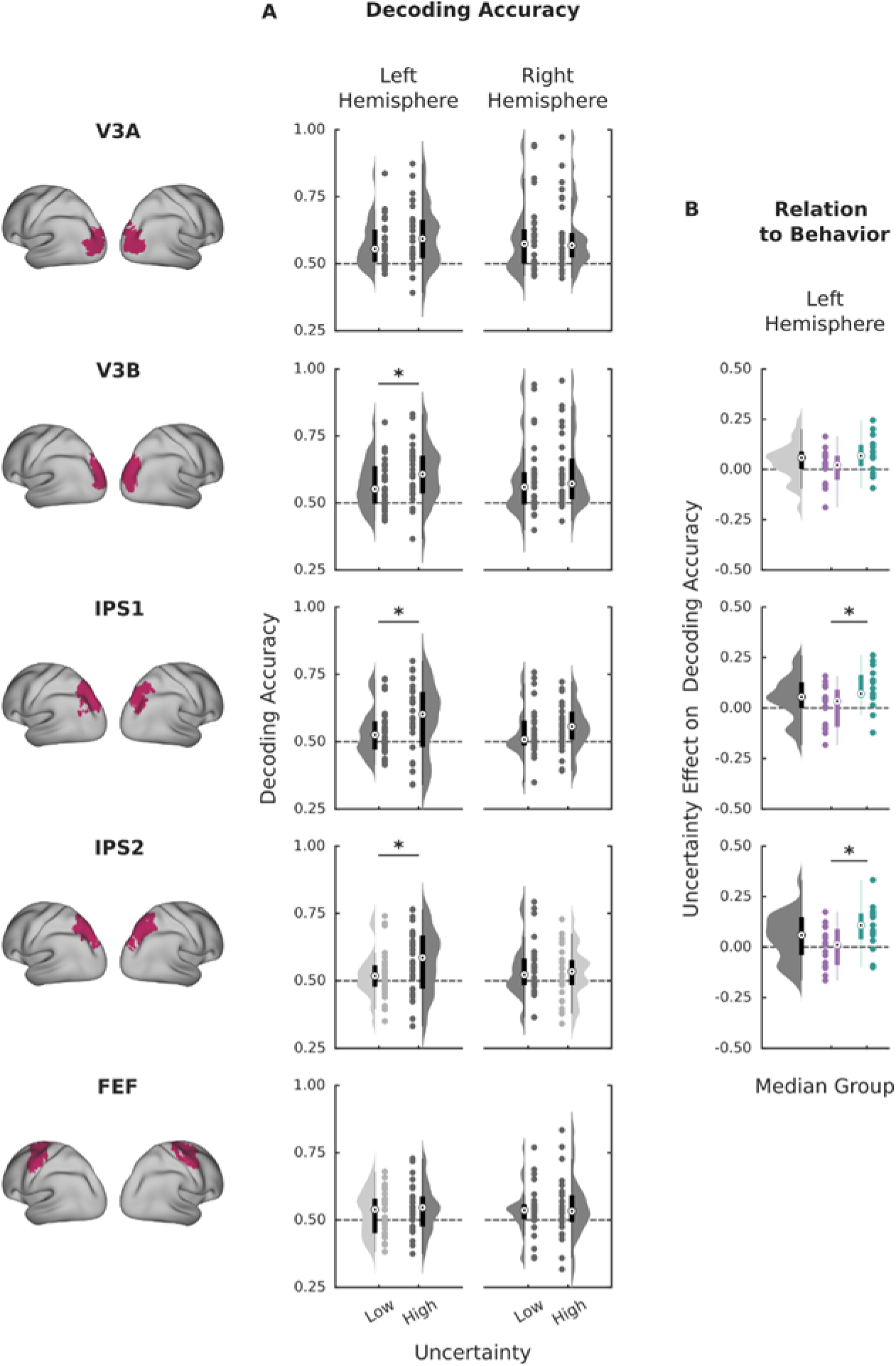
integration of Uncertainty and Diagnosticity in the FP network and relation with attention. **A. Decoding accuracy for the hiD location**. For each ROI, each point is the average accuracy in classifying trials according to whether the hiD location was in the contralateral vs ipsilateral hemifield, for trials with low and high uncertainty. Each point is one participant, and the box plots show the range from the 25^th^ to 75^th^ percentiles, and the median value (white dot). The half violin plot shows the probability density. Dark gray shading indicates ROIs in which decoding was significantly above chance (p < 0.05 vs shuffled controls). Stars indicate significant differences between high and low uncertainty trials (p< 0.05, Wilcoxon paired test). **B.** Uncertainty effects in behavior and neural responses. For each participant and each ROI, we calculated the difference in decoding accuracy between high uncertainty and low uncertainty trials. The box plots and violin plots on the left of each panel show the distributions of these differences across all participants (N = 30) using the same format as in **A**. The colored box plots and points to the right of each panel show the same data for participant groups with attention interaction indices above and below median (defined in Fig. 1C). Stars show p < 0.05, unpaired test between the two participant groups.

In the critical analysis, we recomputed the decoding accuracies for the hiD location for each level of uncertainty. We reasoned that areas that mediate EIG-based control should signal the hiD location with higher accuracy on hiU versus loU trials. Consistent with this hypothesis, the accuracy in decoding the hiD location was significantly enhanced by uncertainty in areas V3B, IPS1 and IPS2 in the left hemisphere (Fig. 3B; paired t-test, hiU vs loU: left V3B (t(29)=2.5, p = 0.0174, CI = [0 0.08]); left IPS1 (t(29)=2.5, p= 0.0174, CI = [0 0.09]); left IPS2 (t(29)=2.3, p = 0.028, CI = [0 0.09]); N = 30 participants). We next tested if these uncertainty modulations differ according to the uncertainty effects on d’ (Fig. 1D). Indeed, the uncertainty effects on spatial decoding accuracy in left IPS1/2 were stronger in participants with above vs below-median d’ interactions (Fig 3B, unpaired t-tests: left IPS1: t(28)=2.3, p= 0.0269, CI = [0.02 0.16]; left IPS2, t(28)=2.5, p= 0.0174, CI = [0.02 0.18]; N = 15 in each group). Thus, portions of the fronto-parietal network integrate diagnosticity and uncertainty consistent with EIG-based attention control.

An important concern is whether these findings reflect effects on arousal or decision formation rather than selective attention. Indeed, as may be expected from prior reports^14,15^ higher uncertainty was associated with larger pupil diameter before and after 2^nd^ letter onset (Fig. S1A-C). Moreover, as expected from the fact that participants integrated evidence in near-Bayesian fashion, uncertainty and diagnosticity affected the decision weight of the 2^nd^ letter (the extent to which this letter’s orientation influenced the final decision about the jar type; Fig. S1D-E). However, extensive analyses showed that task effects differed across behavioral metrics (Fig. S1A-F). Crucially, the effects of diagnosticity and uncertainty in pupil size, decision weights and d’ were uncorrelated (Fig. 2E) and no differences in spatial decoding accuracy were found when participants were split according to the interaction indices in pupil size and decision weights rather than selective attention (2-way ANOVA confirmed an effect of group (above/below median) for splits based on d’ interactions (post-hoc comparisons, p = 0.03 for IPS1 and IPS2) but no significant differences for other splits, all p’s > 0.05). Thus, fronto-parietal areas are sensitive to uncertainty-diagnosticity interactions in d’ but not pupil size or decision formation.

### Uncertainty modulates interactions with the mPFC

While visual and fronto-parietal areas convey spatially tuned visual information, uncertainty is a domain-general signal encoded in non-visual structures^16^. An important question, therefore, is which areas and pathways convey uncertainty to the fronto-parietal network. Based on its involvement in executive function and responsiveness to uncertainty, the medial prefrontal cortex (mPFC) is a prime candidate for fulfilling this function^11,12,17^ but evidence linking this area to perceptual discriminability is lacking.

To examine this question, we again selected an unbiased mPFC mask based on the literature^18^ (Fig 4A, green) and identified a cluster within this ROI that was significantly modulated by uncertainty (Methods: Ucertainty-GLM; Fig 4A, red; MNI peak coordinates = [- 5 6 57], cluster size = 1712 voxels, p = 0.025, small-volume cluster-level corrected). To understand how this cluster interacts with the frontoparietal ROIs, we conducted a whole-brain psychophysiological interaction (PPI) analysis with the uncertainty cluster as seed and uncertainty as the psychological variable. This identified a single cluster in the left hemisphere that overlapped with area V3A-B (Fig. 4B left panels; MNI peak coordinates = [-29 -76 13], cluster size = 2871 voxels, p = 0.015, cluster-level corrected), whose connectivity with the mPFC uncertainty cluster was stronger for high vs low uncertainty (Fig. 4B, right panel; one-sample t-test: t(29)=5.0, p < 10-3, CI = [0.23 0.55]; although note that the effect size is inflated). Thus, one way in which uncertainty signals may reach the frontoparietal network is through functional coupling between the mPFC and area V3 in the left hemisphere.

**Figure 4:**
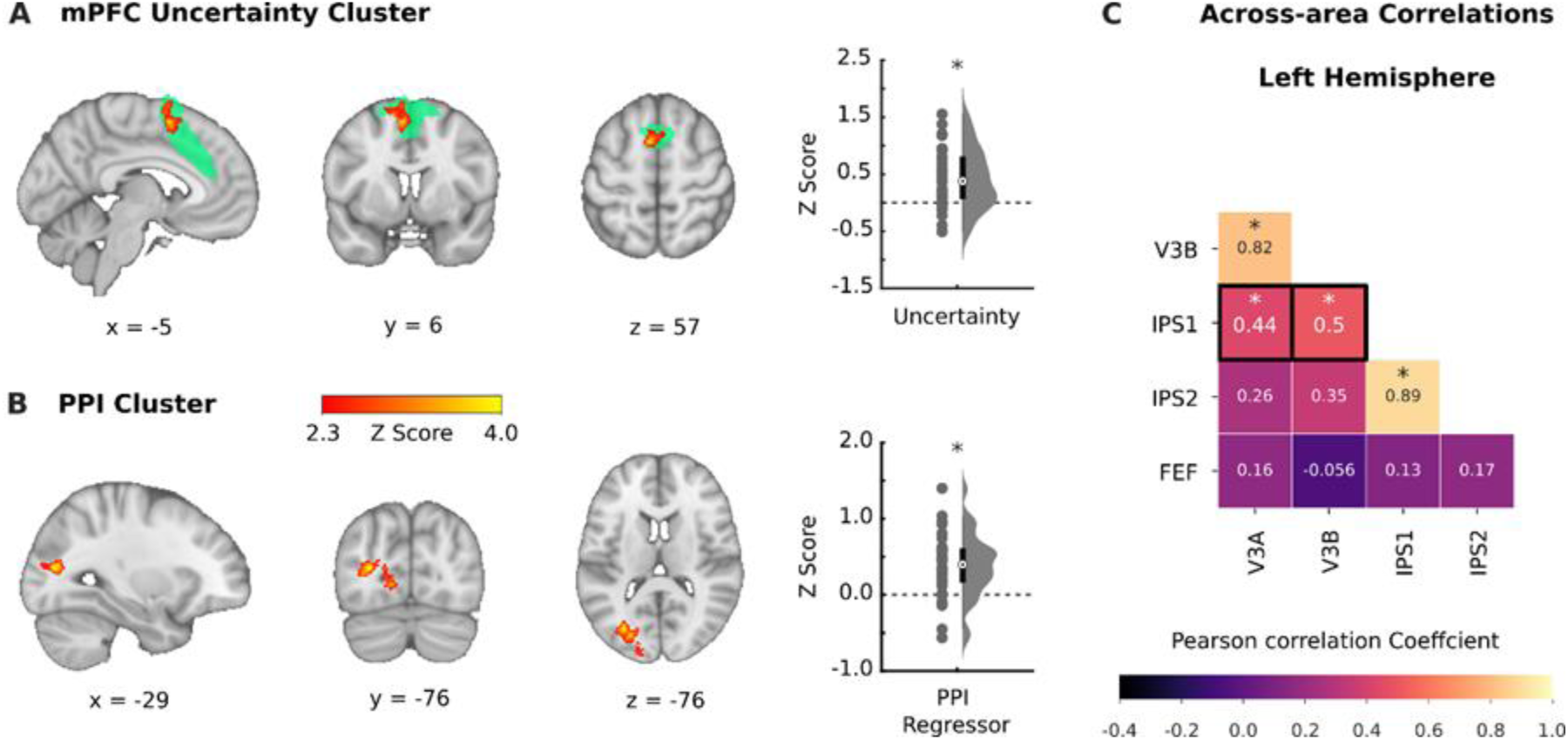
Uncertainty signal involves functional interactions with the executive network. **A. mPFC cluster** encoding uncertainty, including anatomical views of the mPFC mask (green) and the uncertainty-coding cluster inside it (red). The right panel shows the distribution of uncertainty z-scores over the 30 participants, in the same format as Fig. 3A. **B.** Cluster showing significant psychophysiological interactions (PPI) between mPFC and uncertainty, in the same format as **A**. The color bar for z-scores applies to both **A** and **B**. **C. Correlations between the uncertainty modulations of multivariate spatial decoding**, across participants for each pair of uncertainty-sensitive visual and fronto-parietal areas. The colors and numbers show the Pearson correlation coefficient and * denote p < 0.05. The black frames highlight significant correlations between V3 and IPS.

To further track the propagation of uncertainty signals within the fronto-parietal network, we computed the across-subject correlations between the effects of uncertainty on spatial decoding accuracy for each pair of fronto-parietal ROIs. Fig. 4C shows the pattern of these correlations – i.e., the extent to which participants who showed stronger uncertainty effects in one of the posterior ROIs tended to also show stronger effects in other ROIs. Among the uncertainty-sensitive ROIs in the left hemisphere, significant correlations were found between V3A and B and between IPS1 and IPS2 and, more interestingly, between areas V3A and V3B and IPS1 (Fig. 4C, black frames). Note that uncertainty effects were defined as differences in spatial decoding accuracy on high vs low uncertainty trials, and thus this analysis identifies areas that share uncertainty information rather than non-specific BOLD activity. Together with the previous findings, these findings suggest that uncertainty modulations originate at least in part in the mPFC, and are coordinated in a network of posterior fronto-parietal ROIs.

## DISCUSSION

Understanding how attention is deployed as a function of behavioral context has been an enduring challenge in attention research. In traditional tasks of attention, participants are instructed about when and what to attend – e.g., by flashing a salient cue at the to-be-attended location or presenting a symbolic cue that indirectly identifies that location^19^. Although symbolic cues are referred to as “endogenous”, they direct attention by means of an external instruction, independently of the participant’s judgment of the situation and goal^8^. Several recent experiments adopted an alternative approach in which participants are allowed to direct attention at will, and have shown that instructed and willed attention involve distinct interactions between fronto-parietal, cingulo-opercular and dorsal attention networks^20^. However, in these studies the task is held constant, so that attentional fluctuations are purely internally driven and appear random from the experimenter’s standpoint. Here we took a different approach, in which we allowed participants to determine when and where to attend and provided them with a well-defined context in which they could estimate the informativeness of alternative stimuli.

We show that participants allocated covert attention (measured as perceptual discriminability) by integrating diagnosticity and prior uncertainty, approximately consistent with Bayesian EIG. Moreover, this integration was encoded in the fronto-parietal network. Areas V3A/B and IPS1/2 in the left hemisphere provided information about the location of a high versus low diagnosticity stimulus, which became more precise under high versus low prior uncertainty, correlating with the behavioral effects of uncertainty on d’. Connectivity and PPI analyses suggested that uncertainty signals are conveyed through functional interactions between the mPFC and area V3, and propagate through connectivity between V3 and IPS. The results suggest that EIG-based attention control is fundamental not only for overt attention (saccades)^21^ but also covert changes in perceptual discriminability, and reveal the neural mechanisms of this form of control.

### Allocating attention based on uncertainty and diagnosticity

Our manipulation of stimulus diagnosticity is closely related to the construct of predictive validity that is used in prior studies of selective attention^22^. Diagnosticity measures the predictive accuracy of an information source that can emit multiple stimuli (in our experiment, a box from which sample letters were drawn), while validity measures the predictive accuracy of an individual stimulus (in our experiment, an upright or inverted letter “T”). However, tasks that hold uncertainty constant and manipulate only predictive accuracy (whether diagnosticity or validity) cannot fully specify EIG and introduce value confounds, as predictive accuracy correlates with EV^23^.

In the present experiment, we addressed these difficulties by simultaneously manipulating diagnosticity and uncertainty, using a sequential accumulation scenario in which uncertainty changed dynamically as evidence accumulated preceding a choice. We show that, even while maintaining fixation, participants rapidly adjusted their perceptual sensitivity for an anticipated stimulus based on both diagnosticity and uncertainty. These adjustments were distinct from a trial’s EV, and instead were consistent with informational gains -- the marginal decrease in decision uncertainty, relative to prior beliefs, that could be expected from discriminating an additional stimulus.

A salient aspect of our behavioral findings was that, while diagnosticity had a robust effect on d’, the effects of uncertainty showed marked individual variability. The variability in d’ modulations contrasted with the consistent effects of uncertainty on pupil size and decisions, suggesting that, rather than differing in their general sensitivity to uncertainty, participants differed specifically in how they used uncertainty to control perceptual discriminability.

An explanation for this intriguing result may be found in the distinct roles that diagnosticity and uncertainty play in attention control. Whereas diagnosticity describes an information source and specifies where (or to what) to attend, uncertainty specifies when to focus attention. To adopt an uncertainty-sensitive strategy, therefore, our participants had to flexibly adjust their attention in time – that is, by adopting a stronger attentional bias toward the hiD location under higher uncertainty and relaxing this bias in trials with lower uncertainty. However, such a flexible, fine-grained form of control was likely to entail cognitive load. This may have motivated some participants to adopt a simpler strategy of always attending to the hiD location, especially since this strategy had little downside in our task (as the EIG of the hiD location was slightly higher than that of the competing location even on low-uncertainty trials when EIG was overall low). Thus, an interesting hypothesis for future research is that uncertainty sensitivity will become more consistent in tasks that impose higher attentional load and, thus, higher opportunity costs for attending to low EIG stimuli.

Importantly, our results show that the individual variability in diagnosticity-uncertainty integration was encoded in the fronto-parietal network. During the inter-letter delay, multi-voxel activation patterns in left areas V3B and IPS1/2 signaled the location where a hiD 2^nd^ letter would flash with an accuracy that was enhanced by uncertainty and, in left IPS1/2 the uncertainty effects on decoding accuracy corresponded with the uncertainty effects on d’. An important question is whether these areas merely report the selection of an attentional locus fait accompli or, alternatively, integrate diagnosticity and uncertainty for the purpose of selecting this locus. While definitively addressing this question will require a causal manipulation approach, existing results -- on the role of the human fronto-parietal network in advance attentional orienting^4-7^ and neural responses in monkeys -- support the latter interpretation. Consistent with our results, neurons in the monkey lateral intraparietal area, a putative homologue of human IPS, encode diagnosticity^23^ and decision uncertainty in tasks involving saccades^24,25^. Crucially, the neurons encode diagnosticity before signaling the direction of the monkey’s saccade, suggesting that they track the dynamics of the internal selection in advance of an overt motor plan^23^. The hypothesis that the human fronto-parietal network integrates diagnosticity and uncertainty for prioritizing competing locations can thus be tested in future experiments using causal manipulations or higher temporal-resolution recording techniques like EEG or MEG.

Our finding that uncertainty modulated spatial decoding accuracy and multivoxel activity patterns is consistent with the cellular mechanisms by which uncertainty may modulate visuo-spatial selectivity. Converging evidence shows that uncertainty is conveyed by neurons outside the visual system that lack tuning to visual features (e.g., ^16,26^). These neurons provide global modulations and to topographically-organized visual maps and produce variable, cell-specific effects^24,25^ that would manifest as changes in the multivoxel BOLD pattern. A previous study has shown that norepinephrine, one neuromodulator that is likely to signal uncertainty^27^, enhances visual gains^28^. Our findings go further by suggesting that uncertainty enhances the competition among alternative stimuli (which, in turn, can be mediated by higher visual gains^29^). Thus, uncertainty may be viewed as a global (non-spatial) task feature that changes the activity pattern in visuo-spatial structures^30-33^ to enhance the competitive weight of high diagnosticity stimuli and filter out less diagnostic distractors, at the precise times when information is most needed in a particular task.

A notable aspect of our results is that, while fronto-parietal ROIs encoded diagnosticity-uncertainty integration for d’, they did not encode the analogous integration for the final decision regarding jar type. After discriminating the 2^nd^ letter orientation, participants integrated the evidence from the two letters consistent with Bayesian rules, affording more weight to the 2^nd^ letter when it had higher diagnosticity and resolved more prior uncertainty. However, the effects of diagnosticity and uncertainty on d’ and decision formation were uncorrelated, suggesting that Bayesian integration for the two processes rely on distinct mechanisms. Thus, individuals who, in their final decision, rely on sensory evidence more heavily in settings with higher diagnosticity and uncertainty, do not necessarily discriminate the sensory evidence more efficiently in these settings. Moreover, fronto-parietal areas are specifically implicated in the EIG-based modulation of sensory discriminability, while Bayesian integration for the final decision relies on distinct mechanisms.

Our finding that the mPFC is a source of uncertainty signals to the fronto-parietal network is consistent with the role of this area in exploration and executive function^10,34^. However, direct evidence linking the mPFC with perceptual discriminability is lacking. Our connectivity analyses suggest that uncertainty signals from the mPFC reach posterior portions of fronto-parietal network, providing direct evidence that this area signals uncertainty for the purpose of controlling attention and perceptual discriminability^12^.

### Limitations and future directions

The effects of uncertainty we report are more spatially localized than those in previous studies, a finding that is most likely explained by the specific effects of uncertainty in our task. A recent study describes widespread activation throughout frontal areas and the locus coeruleus in response to arousal-inducing fearful stimuli35, while other studies report activations throughout and beyond executive and fronto-parietal areas in tasks in which the uncertainty about an attentional target was manipulated in blocks^36,37^. In contrast, the changes in uncertainty in our task were transient and short-lived, during the anticipatory delay in each trial. Moreover, although these changes produced pupil size modulations, we focused on their specific and independent effects on d’. Thus, the circumscribed activation we find reflects the specific effects of uncertainty on d’ rather than generalized arousal reactions.

The lack of significant uncertainty modulations in the right fronto-parietal ROIs or in the FEF may appear surprising, especially in view of the anatomical proximity of the FEF to the mPFC. However, these negative findings must be interpreted with caution, especially given the variable effects of uncertainty on d’ in our task. Thus, further studies are needed to ascertain whether and how the right versus left hemispheres, and frontal versus visual and parietal areas contribute to uncertainty-based attention control.

In sum, we show that people modulate their perceptual discriminability consistent with information gains when accumulating evidence relevant to a choice, and this process is mediated by the fronto-parietal and executive networks.

## AUTHOR CONTRIBUTIONS

F.J.D.Z. conceptualization, methodology, software, validation, formal analysis, investigation, data curation, writing-original draft, writing-review & editing, visualization. G.H. methodology, resources, writing-review & editing, supervision. J.P.G. conceptualization, methodology, formal analysis, resources, writing-original draft, writing-review & editing, visualization, supervision, funding acquisition.

## ACKNOWLEDGEMENTS

The authors would like to thank members of the Gottlieb lab for their valuable insights. They would also like to thank Ray F. Lee for technical support with the MRI scanner. This work was supported by a Human Frontiers Research Grant to J.P.G.

## METHODS

### Participants

All experimental procedures were approved by the Institutional Review Board of Columbia University. Thirty individuals (20 female) were recruited via mailing lists and a research participation pool at Columbia University, gave written informed consent, and completed the study in the laboratory for monetary compensation (see below). All participants were right-handed with normal or corrected-to-normal vision and no self-reported past or present neurological conditions, were under 40 years of age (range [18–39], mean = 25 years) and passed a health and safety screening on their eligibility for the MRI scanner.

### Task (scan session)

During the scan session, participants viewed visual stimuli that were presented by a projector located behind the scanner bore and reflected through an angled mirror, and responses were recorded via an MR-compatible button-box. The Psychophysics Toolbox for MATLAB (Psychtoolbox) was used for all aspects of experimental control, and infrared video-based eye trackers (EyeLink 1000, SR Research) for recording eye position and pupil size (1,000 Hz sampling rate, monocular tracking).

Each trial of the full task began with the onset of a central fixation cross and the requirement that the participant fixates the cross for 500 ms. Afterwards, the 1^st^ letter was shown for 1 second superimposed on the fixation cross and surrounded by a frame indicating diagnosticity. Two peripheral frames centered at 7° to the right and left of fixation then appeared indicating the possible locations where the 2^nd^ letter could flash. After a variable anticipatory period of 4-9 seconds (drawn randomly from a gamma distribution) the 2^nd^ letter flashed briefly in one of the frames. After the flash, participants indicated if the letter was upright or inverted and then reported their final decision about the majority orientation in the jar (in both cases by pressing the up/down arrows on the keyboard). At the time of the final decision, participants saw a “reminder” screen depicting the two letters they saw surrounded by their respective frames. The 1^st^ letter was on the left and was shown in its veridical orientation (upright or inverted); the 2^nd^ letter was on the right and had the orientation that the participant reported. This procedure minimized memory confounds while ensuring that sensory accuracy was consequential for decision accuracy. After reporting the jar type and receiving feedback about the accuracy of this choice, participants waited for a variable intertrial interval (4-9 seconds) and proceeded to the following trial. If the participant broke fixation at any point before the 2^nd^ letter flash, a message saying “please keep looking at the fixation cross” appeared and the trial was added to the sequence to be repeated later.

The identity of the jar type (majority upright or inverted), and the 1^st^ and 2^nd^ letter diagnosticities were independently randomized from a 50:50 uniform prior. High and low diagnosticity were indicated by red/blue colors, with the color-diagnosticity mapping counterbalanced across participants. The stimuli indicating diagnosticity were square frames subtending 1.3° of visual angle. The fixation cross and letters were black on a gray background and subtended 1° of visual angle. The 2^nd^ letter was preceded and followed by a black grid mask, and was accompanied by a distractor (a different grid pattern) in the opposite frame. The flash duration for the 2^nd^ letter was customized for each participant during the prescan session to support ∼75% discrimination performance (see below, Attention task).

During MRI scanning, participants performed 4 runs of 40 trials each, for a total of 160 trials. To minimize the need for frequent changes in attentional strategies (which could create cognitive load), we imposed additional structure in each run. Each run consisted of two sets of 20 trials with a fixed hiD/loD geometry (the hiD location being in the right/left hemifield), and each geometry set consisted of two sets of 10 trials with a fixed 1^st^ letter diagnosticity. Thus, prior uncertainty was fixed across 10 consecutive trials and the hiD location was fixed across 20 consecutive trials.

At the end of the session, one trial was randomly selected from those that the participants played. Participants received a $1 bonus if they correctly reported the jar type on the selected trial plus a base pay of $20 per hour for participation.

### Pre-Scan session

Before performing the task in the scanner as noted above, all participants completed a pre-scan session that served familiarize them with the task and obtain perceptual thresholds. The session was conducted on a separate day in a testing room outside the scanner equipped with an EyeLink 1000 eye tracker.

During the pre-scan session, participants completed two simplified variants of the task. Participants started with the Inference Task, which introduced the jar types, diagnosticity manipulations and posterior probabilities associated with the 1^st^ and 2^nd^ letter combinations. Participants first received written instructions about the task, the hidden jar and diagnosticity types, followed by comprehension quizzes and 2-3 practice trials, and, finally, 96 trials of the inference task. Similar to the main task, task participants saw two letters on each trial that were surrounded by colored frames indicating diagnosticity. Unlike the main task, both letters were shown clearly at the center of the screen and remained visible throughout the participants’ report. Rather than reporting the 2^nd^ letter orientation, participants reported the posterior probabilities of a jar type, by dragging a mouse on a scale bar ranging from 0% to 100% probability of a majority upright jar. Participants made this report after seeing each letter and received feedback in the form of a red line indicating the correct probability at each step. Finally, participants reported their final decision about the jar type and received feedback on whether their guess was correct.

The task provided participants with the opportunity to learn the posterior probabilities associated with the 16 possible combinations of diagnosticity and orientation using direct feedback. We reasoned that participants who did not learn these posteriors could not be expected to compute EIG or orient attention accordingly. Thus, participants were invited for the following scan session if, by the end of this task, reported posterior estimates within 10% of the Bayesian estimates.

After the inference task, participants performed the Quest and Full Tasks under eye tracking. For this, participants placed their head on a chinrest at 60 cm from the screen and completed an initial 13-point calibration routine. Pupil and corneal reflection detection parameters were continuously monitored and adjusted, and the calibration routine was repeated, if necessary, during the session. The Quest task familiarized participants with the near-threshold discrimination procedure and selected the flash duration that would be used in the full task. Participants viewed a screen containing a fixation cross and two peripheral frames, which was visually identical to the anticipation screen they would see in the full task except that the frames did not convey diagnosticity (were black rather than colored). After maintaining central fixation for 500 ms, participants saw a brief flash of an upright or inverted letter T in one of the frames and reported its orientation by pressing the up or down arrows on the keyboard. Participants completed two blocks of 60 trials in which the location of the “T” was constant (and cued in advance) in the right or left hemifields, and the letter’s duration was adjusted using a QUEST adaptive staircase procedure to 75% correct performance in each hemifield^38^. We then chose the longer of the two threshold durations as the flash duration used in the main task which, across participants, ranged between 15-564 ms (mean=90, SD=10).

Finally, participants performed 160 trials of the full task, identical to the task they will perform in the scanner (previous section and Fig. 1B). At the end of the pre-scan session, participants received compensation of $20/hour plus $1 monetary bonus if their discrimination accuracy was correct on a random trial from those they performed on each of the Inference, Quest and full task blocks (for a maximum possible bonus of $3).

### MRI data acquisition

fMRI images were collected on a 3T Siemens Magnetom Prisma scanner with a 64-channel head coil. The scanner pulse timing was synchronized with the task using the Psychophysics Toolbox for MATLAB (Psychtoolbox). Functional images from 4 runs were obtained with a multiband echo-planar image (EPI) sequence (repetition time = 2s, echo time = 30 ms, flip angle = 80°, acceleration factor = 3, voxel size = 2 mm isotropic; phase encoding direction: posterior to anterior), with 69 axial slices (14° transverse to coronal) acquired in an interleaved fashion. Whole brain high resolution (1.0 mm isotropic) T1-weighted structural images were acquired with a magnetization-prepared rapid acquisition gradient-echo (MRPRAGE) sequence. Finally, to regress out the effect of physiological noise, we collected cardiac and respiratory data using pulse oximetry and a respiratory cushion.

### Data Analysis

#### Behavior

All behavioral data were extracted and analyzed using custom-made Matlab and Python scripts. For each combination of diagnosticity and uncertainty, we computed d’ and criterion^39^. Statistical comparisons were made with t-tests (paired or unpaired, as appropriate) and evaluated at p < 0.05 two-tailed unless otherwise stated. Additional analyses of pupil size and the final decision are described in Supplementary Note 1.

#### Simulations

We simulated an agent who derives the Bayesian posterior probability after observing two random sequential samples and decides on the jar type using a maximizing strategy (i.e., choosing the jar type that corresponds with the higher probability, and choosing randomly for P = 0.5). To simulate the effect of attention, we assumed that the agent has variable probabilities of correctly discriminating the 2^nd^ letter orientation (0.5 to 1 in 0.1 increments). We repeated the simulations 30 times for each discrimination accuracy, emulating the number of trials and trial structure in the real experiment. We used the results to calculate EIG as the slopes of decision accuracy versus perceptual accuracy (Fig. 2A), and expected value (EV) as the decision accuracy at perfect perceptual accuracy (Fig. 2B).

#### fMRI preprocessing

We pre-processed MRI data using FMRIB Software Library (FSL) version 6.0^40,41^ MPRAGE anatomic images were skull stripped using BET. For each run, EPI functional images were motion correction using MCFLIRT in FEAT, high-pass filtered (cut-off = 100 ms), and spatial smoothed (3mm FWHM Gaussian kernel). Functional images were registered to the MPRAGE anatomical image and normalized to the Montreal Neurological Institute (MNI) space using a non-linear transformation with 12 degrees of freedom.

#### Visual and fronto-parietal network analysis

Using the pre-processed data of each participant, we defined functional bilateral FP ROIs for V3A, V3B, IPS1, IPS2, and FEF for each participant using probabilistic maps from Wang et al. 2015^13^.

Within each ROI, we then pooled all the trials and identified the voxels responding during the anticipation epoch using a simple general linear model (GLM). The model included a boxcar function identifying the anticipation period, as well as 6 head-motion regressors estimated by MCFLIRT in the pre-processing stage and 16 voxelwise regressors created by the physiological noise modeling (PNM) tool capturing signal changes due to cardiac and respiratory signals. All regressors were convolved with a double-gamma hemodynamic response function and auto correlations in the time series were corrected with FILM pre-whitening.

For each participant, we then selected the 1000 voxels with the largest beta values in each ROI and used them in the following analyses (similar to ^42,43^). For these responsive voxels, we extracted the responses to uncertainty and diagnosticity during the anticipation period using a standard general linear model. This model (the “Attention GLM”, implemented in FSL FEAT) included 4 regressors of interest – prior uncertainty (low and high) and the hiD location (right and left hemifields) -- alongside the motion regressors described above and nuisance regressors modeling the 2^nd^ letter location (right or left), the 2^nd^ letter orientation (upright or inverted), the perceptual report (upright or inverted) and reaction time (boxcar), the final decision orientation (upright or inverted) and reaction time (boxcar), and the trial feedback (correct or wrong).

We next performed multivariate analyses in which we used linear support vector machine (SVM) classifiers to decode the hiD location (right vs left hemifield) from each ROI on high and low uncertainty trials. We obtained cross-validated accuracy using a leave-one-run-out procedure (training on 3 runs and testing on the 4th) in search light spheres with 3 voxel radius spanning the ROI, extracted the average ROI classification accuracy, and analyzed the results as described in the text. The analyses were implemented in the CoSMoMVPA toolbox^44^ and LIBSVM^45^ for MATLAB.

#### Uncertainty-GLM

We identified the mPFC cluster using a mask from Shirer et al. 2012^18^ We then identified an uncertainty cluster within this mask using the participants’ average beta maps and the Attention GLM to create a group-level uncertainty contrast (high vs low uncertainty) using FSL’s FLAME1. We used a cluster-generating threshold of z > 2.3 and calculated the family-wise-error-corrected p-value at the cluster level based on Random Field Theory, considering a cluster-corrected P<0.05 as statistically significant.

#### PPI Analysis

To perform PPI analysis, we used each participant’s mean BOLD time series across all the voxels of the mPFC uncertainty cluster. We modeled this timeseries using (1) the average BOLD timeseries from other areas (physiological regressors), (2) the onsets of the anticipation delay for high and low uncertainty trials (psychological regressors), (3) the cross-product of the first two regressors (PPI regressor) and (4) all the nuisance regressors from the Attention-GLM. We used participant’s across runs average beta maps to create group-level PPI contrasts using FSL’s FLAME1 and RFT-FWE cluster-level inference as above.

## SUPPLEMENTARY NOTE 1

**Figure S1:**
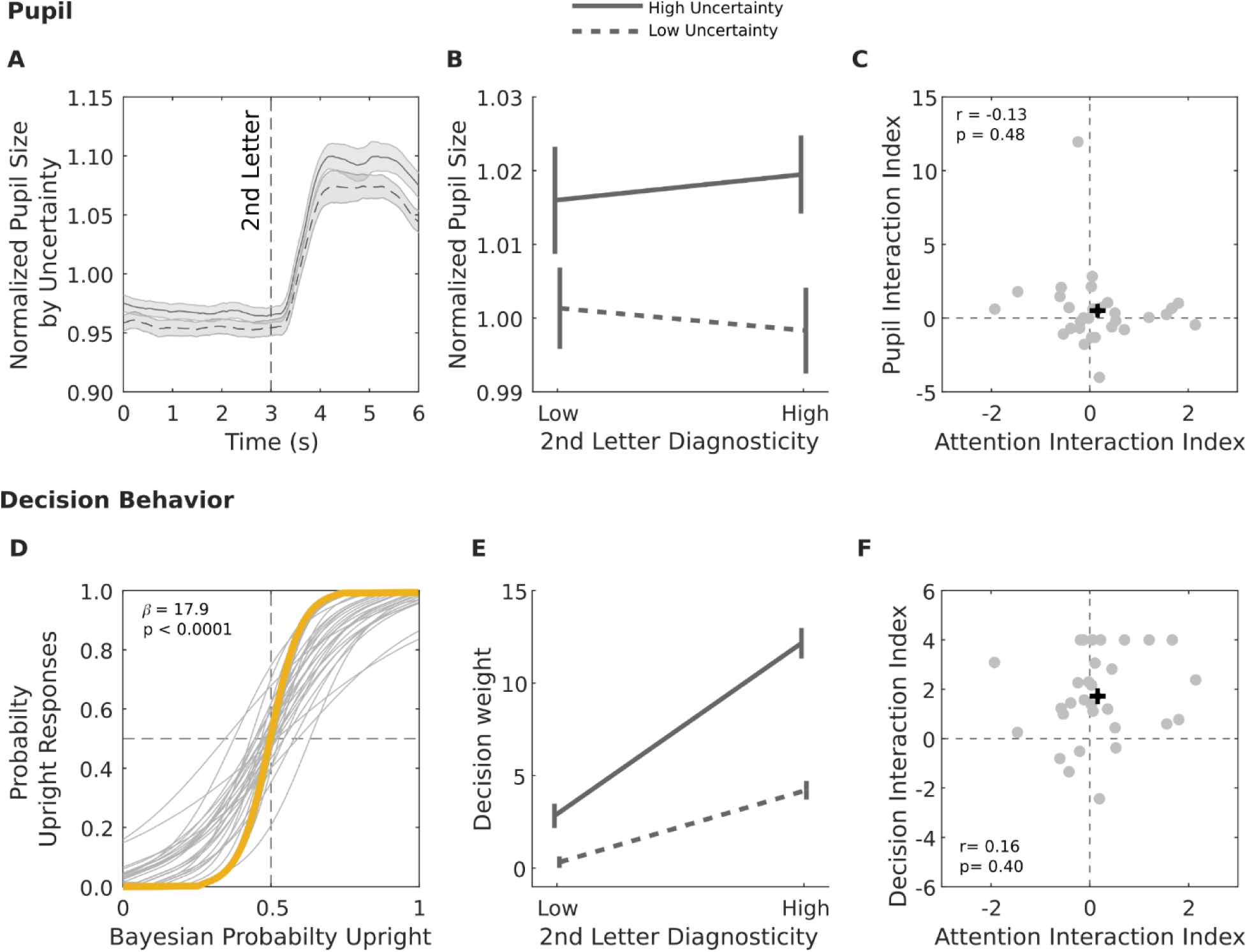
Pupil size and decisions. **A. Pupil diameter** as a function of time relative to 2^nd^ letter onset. For each participant, trials with correct 2^nd^ letter discrimination were averaged for each uncertainty level after pooling across 2^nd^ letter diagnosticities and locations. Traces show average and SEM across all participants (N = 30). **B. Pupil diameter as a function of uncertainty and diagnosticity.** Values show average and SEM across all participants (N = 30) during the 6 second time window in **A**. **C. Interaction indices for pupil vs d’.** Each point is one participant and the black cross shows median values. Dashed lines at 0 indicate no interaction. **C. Decisions versus Bayesian probabilities.** The probability of reporting a majU jar was fit to the Bayesian probability of the jar using mixed-effect logistic regression (see **Supplementary Note 1**). Gray and yellow traces show the fits for, respectively, individual and fixed-effects. **D. Decision weights for the 2^nd^ letter** were fit for each participant and trial type using logistic regression (**Supplementary Note 1**). The points show average and SEM across participants (N = 30). **E. Decision weight and d’ interaction indices are uncorrelated.** All conventions as in panel **C**.

### Pupil size

To pre-process the pupil data, we first identified blinks and saccades events using the Eyelink standard algorithm. For each run, we linearly interpolated blank intervals due to blinks (interpolation time window, from 150 ms before until 150 ms after missing data) and low-pass filtered the resulting pupil time series with a 3^rd^ order Butterworth filter (cut-off: 6 Hz). We then applied least-squares deconvolution to estimate the pupil responses to blink and saccades, and removed them from pupil time series using linear regression^46-48^

To measure pupil diameter, we normalized pupil data to the participant’s mean over the entire session and pooled the normalized values across participants^15^. We focused on a 6 second time window centered on 2^nd^ letter onset (**Fig. S1A**) and also on trials in which participants correctly discriminated the 2^nd^ letter orientation. This strategy provided a sufficiently long averaging window while making it likely that participants correctly registered the 2^nd^ letter diagnosticity after its onset. A 2-way ANOVA showed that pupil size was significantly enhanced by uncertainty (F=8.8, p = p < 10^-5^; N = 30 participants) with no effect of diagnosticity (F=0, p =0.960) or interaction between uncertainty and diagnosticity (F=0.29, p=0.5901).

To compare these effects with those on d’, we calculated diagnosticity, uncertainty and interaction indices for the pupil analogously to those for d’ (*Methods*). Consistent with the ANOVA analysis, the pupil indices for uncertainty, not diagnosticity or interaction, were significantly positive (**Table 1**). Importantly, the indices were not correlated for pupil vs d’ (**Fig. 2E**), as shown for the interaction indices in **Fig. S1C** (r=-0.13, p=0.48, N = 30). Thus, pupil size was affected by uncertainty, consistent with previous work^15,14^, and these effects were distinct from those on d’.

### Decision

We first examined if participants correctly performed the inference task, by plotting their probability of reporting a jar type against the true Bayesian posterior probability (**Fig. S1D**). A mixed effect logistic regression produced a highly significant slope (β = 17.998, p < 10^-3^, CI = [15 21]), confirming that participants used an approximately Bayesian inference strategy.

Next, we examined how much weight participants afforded to the 2^nd^ letter orientation when inferring jar type. To this end, we used lasso regularized logistic regression (the *lassoglm* function in MATLAB) to model, for each participant and each trial type, the trial-by-trial decision about the jar type as a function of the 1^st^ and 2^nd^ letter orientations. This provided us with the decision weight of the 2^nd^ letter orientation after controlling for the 1^st^ letter orientation.

Consistent with the participants’ Bayesian inferences (**Fig. S1A**), a 2-Way ANOVA showed that the 2^nd^ letter weight increased with both diagnosticity (F=45.9, p = 0.965) and uncertainty (F=45.9, p < 10^-5^) with a significant interaction (F=65, p < 10^-5^), a pattern confirmed by index-based analysis (**Table 1**). Importantly, the decision and attention indices were uncorrelated (**Fig. 2E**), and this is illustrated in **Fig. S2C** for the interaction indices (r=0.16, p=0.40, N = 30). Thus, the effects of uncertainty and diagnosticity on d’ could not be explained by the corresponding effects on decisions.

